# Reverse Correlation Uncovers More Complete Tinnitus Spectra

**DOI:** 10.1101/2022.12.23.521795

**Authors:** Alec Hoyland, Nelson V. Barnett, Benjamin W. Roop, Danae Alexandrou, Myah Caplan, Jacob Mills, Benjamin Parrell, Divya A. Chari, Adam C. Lammert

## Abstract

**Goal:** This study validates an approach to characterizing the sounds experienced by tinnitus patients via reverse correlation, with potential for characterizing a wider range of sounds than currently possible.

**Methods:** Ten normal-hearing subjects assessed the subjective similarity of random auditory stimuli and target tinnitus-like sounds (“buzzing” and “roaring”). Reconstructions of the targets were obtained by regressing subject responses on the stimuli, and were compared for accuracy to the frequency spectra of the targets using Pearson’s *r*.

**Results:** Reconstruction accuracy was significantly higher than chance across subjects: buzzing (*M* = 0.53, *SD* = 0.27): *t*(9) = 5.766, *p <* 0.001; roaring (*M* = 0.57, *SD* = 0.30): *t*(9) = 5.76, *p <* 0.001.

**Conclusion:** Reverse correlation can accurately reconstruct nontonal tinnitus-like sounds in normal-hearing subjects, indicating its potential for characterizing the sounds experienced by patients with non-tonal tinnitus.

**Impact Statement:** Characterization of tinnitus sounds can inform treatment by facilitating individualized sound therapies, leading to better outcomes for patients suffering from the cognitive and psychological effects of tinnitus.

## I. Introduction

TINNITUS—the perception of sound in the absence of any corresponding external stimulus—affects up to 50 million people in the U.S. [1], a third of whom experience functional cognitive impairment and diminished quality of life [2], [3]. Clinical guidelines for tinnitus management involve targeted exposure to tinnitus-like sounds in a habituation therapy term *sound therapy*, or as part of cognitive behavioral therapy [4]. Critically, sound therapy outcomes improve when closely informed by the patient’s internal tinnitus experience [5]–[7]. However, existing strategies for characterizing tinnitus sounds, such as *pitch matching* (PM), are best suited for patients whose tinnitus resembles pure tones (*e.g*., ringing), which may represent only 20− 50% of patients [8], [9]. There is a need for methods to characterize nontonal (*e.g*., buzzing, roaring) tinnitus [10], [11].

Nontonal tinnitus sounds are presumed to be complex and heterogeneous [12], although few characteristics have been firmly established. Therefore, we base our approach on *reverse correlation* (RC), an established behavioral method [13]–[15] for estimating internal perceptual representations that is unconstrained by prior knowledge about the representations themselves. RC asks participants to render subjective judgments over random stimuli, and reconstructs the latent signal as a linear combination of stimuli. RC is closely related to Wiener theory, which has inspired “white-noise” approaches to system characterization in physiology [16], [17] and engineering [18].

Here, we validate RC as a method for characterizing a more complete psychoacoustic tinnitus spectrum (PTS) for tinnitus sounds. To that end, normal-hearing participants completed an augmented RC experiment, comparing random stimuli to a target tinnitus-like sound. The estimated PTS was subsequently validated against the targets. Our results demonstrate, for the first time, that tinnitus-like sounds with complex spectra can be accurately estimated using RC.

## II. Materials and Methods

### A. Stimuli

The frequency space of the stimuli was partitioned into *b* = 8 Mel-spaced frequency bins, which divide the frequency space between [100, 13, 000] Hz into contiguous segments of equal amplitude (*i.e*., “rectangular” bins). Reconstruction detail increases with *b*, but *b* = 8 provides a good approximation to the chosen target sounds (*cf*. Fig. 3).

For each stimulus, [2, 7] bins were randomly “filled” with power 0 dB. “Unfilled” bins were assigned − 100 dB. All frequencies were assigned random phase. Inverse Fourier transform of the constructed spectrum yields a 500 ms stimulus waveform.

### B. Target Sounds

Two spectrally complex and complementary target sounds (“buzzing” and “roaring”) were downloaded from the American Tinnitus Association [19] and truncated to 500 ms in duration (Fig. 3).

### C. Experiment

Ten (*n* = 10) normal-hearing subjects listened to *A*− *X* trials containing a target sound (*A*) followed by a stimulus (*X*). *X* wasrandomly generated for each trial, while *A* remained the same within a block of 100 trials (Fig. 1). Subjects completed two (2) blocks per target sound (*p* = 200 total trials per subject). Subjects were told that some stimuli had *A* embedded in them, and were instructed to respond “yes” to such stimuli, otherwise “no.” Subjects listened over earphones at a self-determined comfortable level. Procedures were approved by the UMass IRB.

**Fig. 1.**
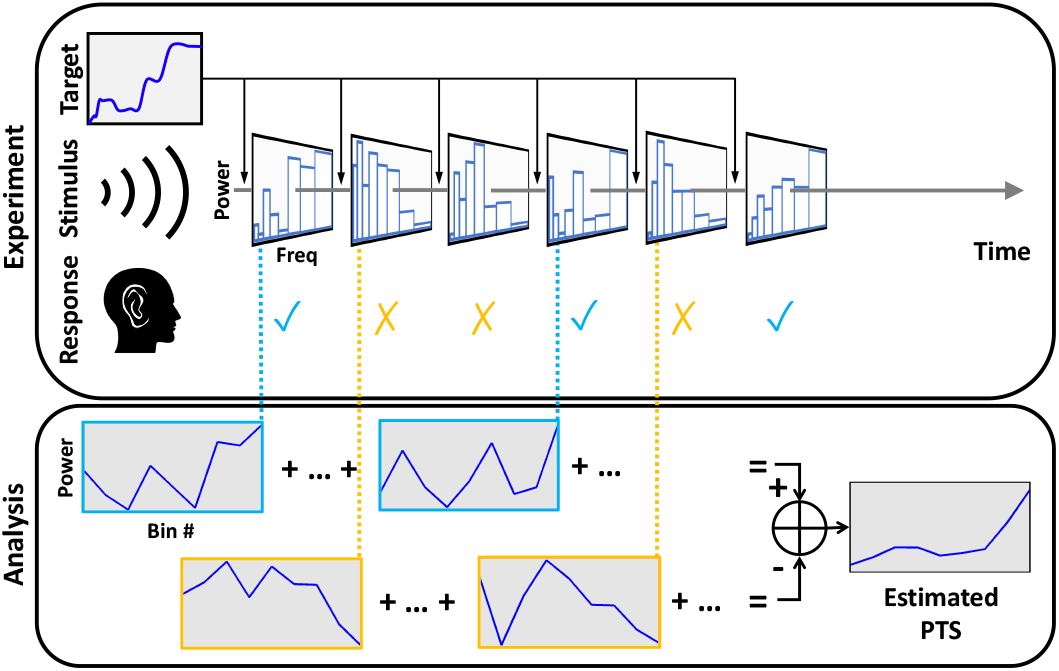
Diagram of the experimental protocol. Subjects listen to a series of random stimuli, each preceded by a target sound. Subjects compare the stimulus to the target, and respond either “yes” or “no” depending on their perceived similarity. The recorded stimulus-response pairs are used to form an estimate of the target.

### D. Reconstruction

A subject performing *p* RC trials with *b* frequency bins produces a stimulus matrix Ψ∈ ℝ^*p*×*b*^ and a response vector *y*∈ { 1, −1} ^*p*^, where 1 corresponds to a “yes” response and− 1to a “no.” RC classically assumes the subject response model:

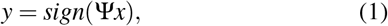

where *x*∈ ℝ^b^ is the subject’s internal representation of interest (*i.e*., of their tinnitus). Inverting this model yields:

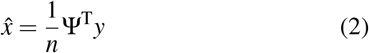

which is a restricted form of the Normal equation under the assumption that the stimulus dimensions are uncorrelated [14].

### E. Validation

The experimental paradigm allows for direct validation of the reconstructions 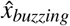and 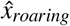 We represent the spectra of the target sounds as vectors *s*_*buzzing*_ ∈ℝ^b^ and *s*_*roaring*_ ∈ℝ^b^ using the same frequency bins as the stimulus with power equal to the mean power at frequencies within that bin. Pearson’s *r* between *s*_*buzzing*_ and *s*_*roaring*_ and their corresponding reconstructions quantifies reconstruction accuracy. One-sample t-tests were performed on the mean Fisher-transformed Pearson’s *r* values across subjects to assess significant differences from zero.

### F. Synthetic Subjects

To establish bounds on human performance, additional experiments were run with two simulated subjects who give either *ideal* or *random* responses. Each experiment ran for *p* = 200 trials and was repeated 1000 times.

The *ideal* subject gives responses following:

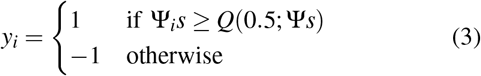

for *i* ∈1, …, *p*, where *Q*(*x, y*) is the quantile function for *x* ∈ [0, 1] of the similarity calculation Ψ*s*, and Ψ_*i*_ is the *i* column of Ψ. Thus, the ideal subject has precise knowledge of every stimulus and responds according to Eq. 3. The *random* subject responds *y*_*i*_∈ {1, − 1} with uniform random probability, thus ignoring the stimulus entirely.

## III. Results

Figure 2 shows the distribution of Pearson’s *r* for human, ideal, and random subject responses. Human accuracies were generally higher than random, with some approaching *ideal*. Mean human accuracy was significantly different from 0 (i.e., mean *random* accuracy) in all conditions: buzzing: *t*(9) = 5.766, *p <* 0.001; roaring: *t*(9) = 5.76, *p <* 0.001; combined: *t*(19) = 7.542, *p <* 0.001. Figure 3 plots the most accurate human reconstructions over the target sound spectra.

**Fig. 2.**
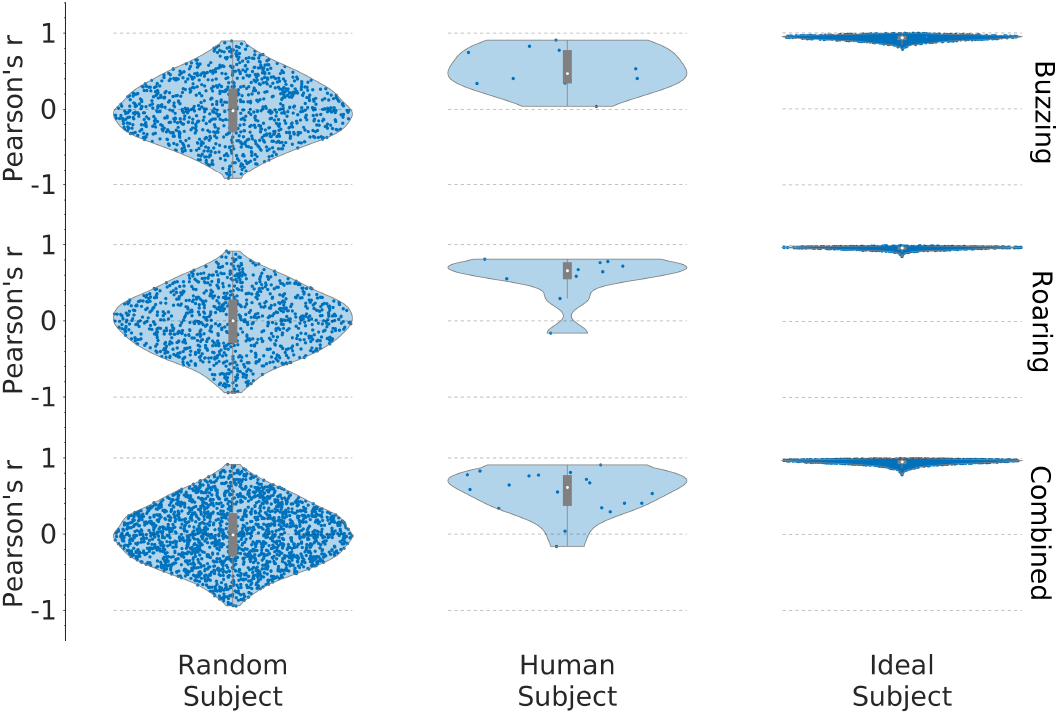
Human reconstruction accuracy is significantly above baseline, but is not optimal. Random, human, and ideal reconstruction accuracies are shown as violin plots with box plots overlaid. The median is a white dot, the ordinate of the blue points are the Pearson’s *r* values.

**Fig. 3.**
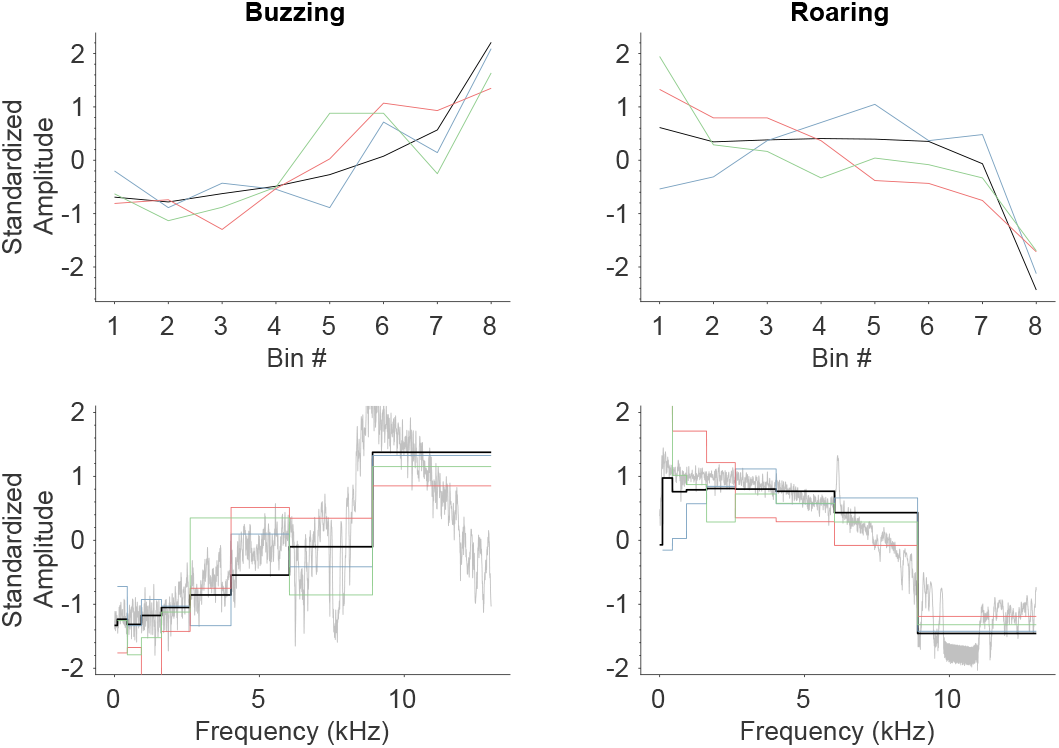
Reconstructions of the PTS capture many salient features of tinnitus spectra. The black trace indicates the target, while colored traces plot exemplar human reconstructions. The top row shows standardized power levels within each frequency bin. The bottom row maps the 8-dimensional bin domain to a 11025-dimensional frequency domain with the unbinned power spectral density of the targets shown in gray.

## IV. Conclusion

Our results show that RC can accurately reconstruct the PTS of complex, non-tonal, tinnitus-like sounds, capturing the most salient features of the target. Reconstruction accuracies observed here are below the simulated ideal, which may be attributed to noisy responses universally observed in applications of RC, and which may be mitigated by further optimizing the experimental protocol, stimulus generation, and reconstruction method. For example, recent approaches to improving RC reconstruction methods can boost efficiency, noise robustness and overall accuracy [20]. Subjects completed the required number of trials within 10 minutes, indicating that this procedure can be conducted in a single clinical visit. RC may therefore be useful as the basis for a clinical assay to characterize a wider variety of tinnitus percepts than currently possible. Future work will focus on validating this approach in patients suffering from tinnitus.

## Footnotes

1 Software for the experiments and analysis was written in MATLAB and is freely available at https://github.com/alec-hoyland/tinnitus-reconstruction/

